# Methylation-sensitive expression of a DNA demethylase gene serves as an epigenetic rheostat

**DOI:** 10.1101/015941

**Authors:** Ben P. Williams, Daniela Pignatta, Steven Henikoff, Mary Gehring

## Abstract

Genomes must balance active suppression of transposable elements (TEs) with the need to maintain gene expression. In Arabidopsis, euchromatic TEs are targeted by RNA-directed DNA methylation (RdDM). Conversely, active DNA demethylation prevents accumulation of methylation at genes proximal to these TEs. It is unknown how a cellular balance between methylation and demethylation activities is achieved. Here we show that both RdDM and DNA demethylation are highly active at a TE proximal to the major DNA demethylase gene *ROS1*. Unexpectedly, and in contrast to most other genomic targets, expression of *ROS1* is promoted by DNA methylation and antagonized by DNA demethylation. We demonstrate that inducing methylation of the *ROS1* proximal region is sufficient to restore *ROS1* expression in an RdDM mutant. Additionally, methylation-sensitive expression of *ROS1* is conserved in other species, suggesting it is adaptive. We propose that the *ROS1* locus functions as an epigenetic rheostat, tuning the level of demethylase activity in response to methylation alterations, thus ensuring epigenomic stability.

**Author Summary:** Organisms must adapt to dynamic and variable internal and external environments. Maintaining homeostasis in core biological processes is crucial to minimizing the deleterious consequences of environmental fluctuations. Genomes are also dynamic and variable, and must be robust against stresses, including the invasion of genomic parasites, such as transposable elements (TEs). In this work we present the discovery of an epigenetic rheostat in plants that maintains homeostasis in levels of DNA methylation. DNA methylation typically silences transcription of TEs. Because there is positive feedback between existing and *de novo* DNA methylation, it is critical that methylation is not allowed to spread and potentially silence transcription of genes. To maintain homeostasis, methylation promotes the production of a demethylase enzyme that removes methylation from gene-proximal regions. The demethylation of genes is therefore always maintained in concert with the levels of methylation suppressing TEs. In addition, this DNA demethylating enzyme also represses its own production in a negative feedback loop. Together, these feedback mechanisms shed new light into how the conflict between gene expression and genome defense is maintained in homeostasis. The presence of this rheostat in multiple species suggests it is an evolutionary conserved adaptation.

## INTRODUCTION

In plants, animals, and fungi, DNA methylation is used to repress the transcription of potentially harmful DNA sequences [1]. Targets include long transposable elements (TEs) that have intact open reading frames and primarily reside in heterochromatin, as well as fragments of TEs that are prevalent in euchromatic gene-rich regions. In plants, DNA methylation is dynamically regulated during development and in response to external perturbations. Many of these changes occur at TEs or TE-derived sequences. Examples include modest DNA methylation changes in gene-proximal regions upon exposure to bacteria or bacterial elicitors [2,3] and DNA demethylation of TEs in the 5′ regions of stress response genes during fungal infection [4]. Dynamic methylation changes have also been implicated in the regulation of genes in response to abiotic signals [5,6]. Furthermore, DNA methylation is dynamic during reproductive development. DNA demethylation in the female gametophyte is important for establishing gene imprinting in the endosperm after fertilization [7]. During male gametogenesis, the sperm become hypomethylated in certain sequence contexts [8]. Similar to other dynamic changes, the removal of methylation in gametophytes occurs largely at TE fragments in euchromatin.

DNA methylation patterns are a product of methylation and demethylation activity, but how these opposing activities are balanced in the genome is unknown. In plants, DNA methylation is established and maintained in different cytosine sequence contexts by genetically distinct pathways. Euchromatic TEs in Arabidopsis and maize are primarily targeted for cytosine methylation through the process of RNA-directed DNA methylation, which results in cytosine methylation in all sequence contexts (CG, CHG, and CHH, where H represents any base other than G) [1,9]. This process is initiated by transcription of non-coding RNAs by a specialized RNA polymerase unique to plants, RNA Pol IV. These non-coding RNAs are then converted into dsRNAs by the RNA-dependent RNA polymerase RDR2. Small 24 nt RNAs generated from these transcripts are then loaded into AGO4. The small RNAs are thought to interact with non-coding transcripts that are generated by a second plant-specific polymerase, RNA Pol V [10], resulting in the recruitment of the *de novo* methyltransferase DRM2 and sequence-specific DNA methylation. 21-22 nt small RNAs generated through an RDR6-dependent pathway can also direct *de novo* methylation independently of RNA Pol IV [11,12]. Positive feedback between existing and *de novo* DNA methylation reinforces silencing [13,14]. Maintenance of asymmetric CHH methylation requires continual *de novo* methylation by RdDM. Other processes maintain DNA methylation in the CG and CHG sequence context. CG DNA methylation is maintained by the maintenance methyltransferase MET1 in conjunction with VIM methyl-binding proteins. CHG methylation is maintained by CMT3 and is positively reinforced by histone H3K9 dimethylation [15]. By contrast, TEs in heterochromatic sequences are methylated by CMT2 with the assistance of the nucleosome remodeler DDM1 [16,17].

Because there is positive feedback between methylation and further RdDM activity and because many RdDM targets are near genes, it is important that mechanisms are in place to protect genes from potentially detrimental hypermethylation. In the Arabidopsis genome, 44% of genes have a TE within 2 kb of the transcribed region [18], potentially creating a conflict between TE suppression by RdDM and gene expression. DNA methylation is opposed by 5-methylcytosine DNA glycosylases that remove methylcytosine from DNA by base excision repair. Plants with mutations in the three DNA glycosylases expressed in somatic tissues, *ROS1, DML2,* and *DML3,* gain methylation in all sequence contexts in gene proximal regions, primarily around TEs and TE-derived sequences [19,20]. DNA demethylation is therefore important to protect genes from RdDM spreading. This has been demonstrated at several loci. For example, the *EPF2* gene, which is associated with a methylated TE approximately 1.5 kb 5′ of the transcriptional start site, gains methylation in the region between the TE and 5′ end of the gene in *ros1 dml2 dml3* mutants, resulting in transcriptional silencing [21]. Although DNA methylation is primarily thought of as repressive to transcription [20,22], expression of the DNA demethylase gene *ROS1* is unexpectedly reduced in some DNA methylation mutants [23–26], although whether this is a direct or indirect effect has not been demonstrated. These observations on *ROS1* expression form the basis of our study.

Here we describe the existence of a rheostat for genomic methylation activity. We find that RdDM and DNA demethylation converge on TE-derived sequences 5′ of *ROS1*. In contrast to other genomic targets of these pathways, expression of *ROS1* is promoted by the RdDM pathway and inhibited by demethylation by ROS1. Thus the *ROS1* locus functions as a self-regulating epigenetic rheostat, balancing input from both DNA methylation and demethylation to maintain homeostasis between these opposing systems.

## RESULTS

### RNA-directed DNA methylation promotes *ROS1* expression

Previous studies have shown that *ROS1* expression is reduced in mutants in which DNA methylation is disrupted or altered [23–26]. Expression of the *ROS1* gene is significantly reduced when plants are grown on the methyltransferase inhibitor 5-aza-2-deoxycytidine (5-azaC) [24] (Fig.1A). Here we systematically evaluated which DNA methylation pathways promote *ROS1* expression by performing RT-qPCR on multiple Arabidopsis mutants that directly or indirectly alter DNA methylation. *met1* plants have pleiotropic methylation phenotypes; methylation in CG, CHG and CHH sequence contexts is reduced genome-wide in combination with local regions of non-CG hypermethylation [20]. We observed extremely low levels of *ROS1* transcripts in *met1* and *vim* seedlings (Fig. 1B), as has been reported previously [23,24]. An approximately ten-fold decrease in *ROS1* transcript levels was observed in eleven different RdDM mutants (Fig. 1C), consistent with previous findings that *ROS1* expression is reduced in *rdr2, nrpd1a, nrpd1b, drd1,* and *drm2* mutants [23,25]. RdDM is predominantly associated with transcriptional repression; therefore transcriptional activation of *ROS1* by RdDM potentially represents an under appreciated function for this pathway. Transcripts from the related 5-methylcytosine DNA glycosylases *DML2* and DML3 are present at much lower levels than *ROS1* in wild-type tissues (S1 Fig.). Mutations in the RdDM pathway do not alter the transcript abundance of *DML2,* and result in small reductions in of *DML3* (S1 Fig.). No significant changes to *ROS1* transcript abundance were observed in CHG methyltransferase mutants (Fig. 1D), in mutants of key regulators of histone H3K9 methylation (Fig. 1D), which is tightly associated with CHG methylation [27], in plants with mutations in genes required to establish non-CG methylation in heterochromatin [16,17] (Fig. 1E), or plants with a mutation in the *RDR6* gene, which can trigger *de novo* methylation independently of the canonical RdDM pathway [11,12] (Fig. 1E). Thus, *ROS1* down-regulation in methylation mutants is restricted to *met1* and its cofactors, and mutants in the RdDM pathway.

**Fig. 1:**
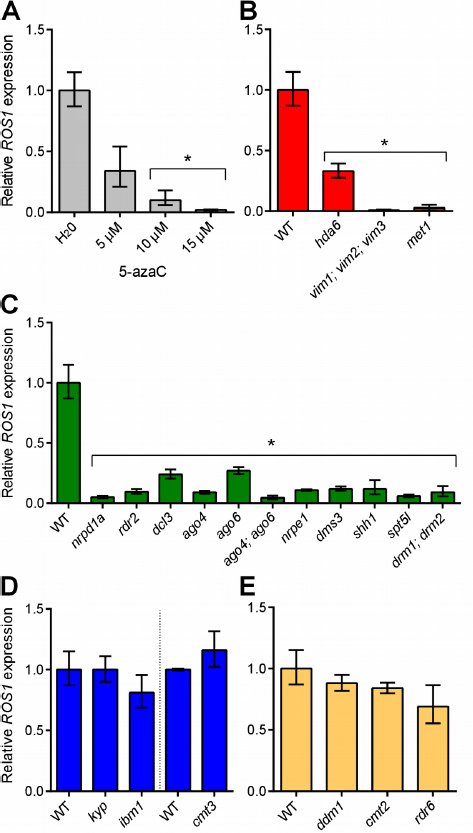
Mutants in the RdDM pathway have reduced *ROS1* expression. **(A)** Relative *ROS1* transcript abundance was measured using RT-qPCR on *A. thaliana* seedlings that were grown on medium supplemented with water or 5 − 15 μM 5-aza-2-deoxycytidine (5-azaC). Relative *ROS1* transcript levels were also quantified in multiple mutants of each of the major DNA methylation pathways in *Arabidopsis* including: **(B)** the maintenance methylation pathway, (C) the RdDM pathway, **(D)** the CHG/H3K9 methylation pathway and **(E)** the heterochromatin DNA methylation pathway or post-transcriptional silencing. WT represents Col-0 in all cases, except for *cmt3, metl* and *drml; drm2,* where the comparative wild-type strains used were Ler, *Col-gl* and Ws-2, respectively. Data represented as mean, error bars represent standard deviation. *p = <0.001, two-tailed t-test.

To test if *ROS1* silencing in *met1* or RdDM mutants was heritable, we crossed *met1, rdr2,* and *drm1 drm2* plants to wild type and evaluated *ROS1* expression in heterozygous F_1_ progeny. *ROS1* expression remained reduced in *MET1/met1* F_1_ progeny at levels about half that of wild type plants, regardless of whether the *met1* plant served as the male or female parent in the cross (Fig. 2A). This suggests that the *ROS1* allele inherited from the *met1* parent remained silenced through meiosis. However, *ROS1* expression was gradually restored as *MET1/met1* progeny developed (Fig. 2B). Thus, erasure of met1-induced epigenetic changes and restoration of normal regulatory mechanisms at *ROS1* likely takes place over multiple cell divisions. By contrast, *ROS1* transcripts were restored to wild-type levels in F_1_ progeny of RdDM mutants crossed to wild type (Fig. 2C).

**Fig. 2:**
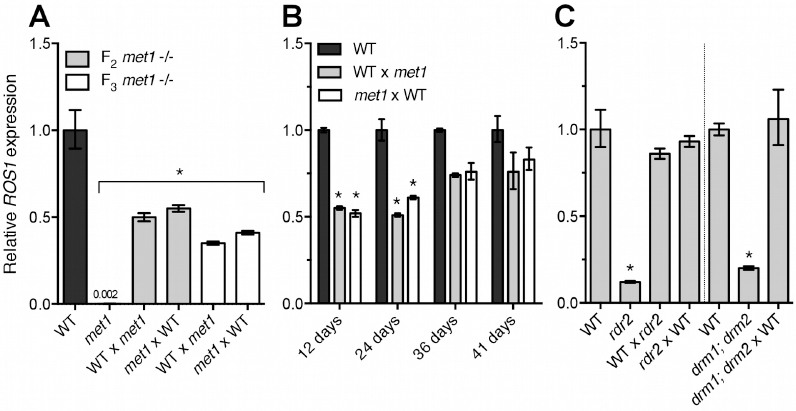
*ROS1* down-regulation in RdDM mutants is not heritable through meiosis. Relative *ROS1* was measured in **(A)** 12 day-old seedlings of F_1_ progeny of homozygous *met1* mutants crossed as either males or females to WT *(met1* parents were either from the F_2_ or F_3_ generation, indicating the first and second generation as homozygous *met1* plants, respectively), **(B)** at different stages during the growth of the *MET1/met1* F1 plants (12 day-old seedlings, 24 day-old rosette leaves and 36 and 41 day-old flower buds) and **(C)** in seedlings of F_1_ progeny of *rdr2* and *drm1; drm2* crossed to WT. WT represents Col-gV, Col-0 and Ws-2 for *met1, rdr2,* and *drm1; drm2* respectively. For crosses, females are written first. Data represented as mean, error bars represent standard deviation. *p = <0.001, two-tailed t-test.

Recently it has been shown that reduced expression of the histone demethylase gene *IBM1* contributes to reduced *ROS1* expression in *met1* mutants [26]. However, we found that *ROS1* expression is not reduced in *ibm1* mutants (Fig. 1D), nor is *IBM1* expression reduced in the RdDM mutants *rdr2* and *nrpd1a* (RNA Pol IV) (S2 Fig.), suggesting that the decreased expression of *ROS1* expression in RdDM mutants is IBM1-independent. Together these data indicate that the down-regulation of *ROS1* observed in *met1* and RdDM mutants are distinct processes, which was further supported by methylation profiling of *ROS1* in different mutant backgrounds, described below.

### RdDM is highly active at a TE 5′ of *ROS1*

To determine if the *ROS1* locus is targeted directly by DNA methylation, we performed bisulfite PCR and sequencing on the entire *ROS1* gene and 1 kb of 5′ flanking sequences (Fig. 3, S3 Fig.). In wild-type plants we identified two small regions where cytosines were methylated in CG, CHG, and CHH sequence contexts (a hallmark of RdDM): a 228 bp region partially overlapping an AtREP5 TE directly upstream of *ROS1,* and in sequences encoding exons 15-18 (Fig. 3). Genome-wide chromatin-IP datasets [10,28] showed that peaks of the RdDM proteins NRPE1 and AGO4 were present 5′ of the *ROS1* start codon, overlapping the nearby TE (Fig. 2). This is the same region where an RNA Pol V transcript has been detected [10]. There were high levels of CHH methylation in this region, predominantly on the top strand (Fig. 2, S3 Fig.), and we identified multiple 24 nucleotide small RNAs directly matching this sequence from published datasets (Fig. 3) [29,30]. In addition, we detected multiple small RNAs matching the methylated exons within the *ROS1* coding region, but these did not overlap with peaks in the AGO4 or NRPE1 ChIP datasets, consistent with the low levels of CHH methylation in this region (Fig. 3). To determine the proximity of the 5′ methylated region to the *ROS1* transcriptional start site (TSS), we performed 5′ RACE using RNA from wild-type Col-0 seedlings and identified two transcription start sites, 26 and 442 bp 5′ of the *ROS1* start codon (S3 Fig.), the latter of which is within 100 bp of the methylated region.

**Fig. 3:**
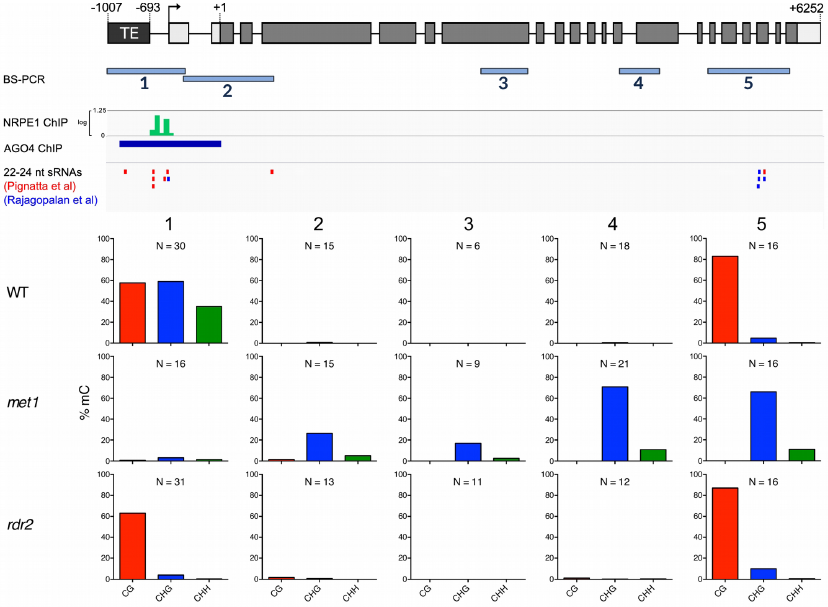
*ROS1* is an RdDM target. Methylation of the entire *ROS1* locus was examined in wild-type by bisulfite sequencing. Bisulfite sequencing from WT (Col-0 and Col-g/), *metl,* and *rdr2* rosette leaves is shown for five regions. The activity of RNA polymerase V (NRPE1-ChIP [28]), and AGO4 (AGO4-ChIP [10]) is shown in tracks below the *ROS1* locus. 22-24 nucleotide small RNAs matching *ROS1* were identified from two published datasets [29,30]. N represents the number of independent clones sequenced for each bisulfite PCR product.

We also profiled *ROS1* methylation in *met1* and *rdr2* mutants. CG methylation was eliminated in *met1,* but the *ROS1* coding region was hypermethylated in the CHG context (Fig. 2), as has been previously reported [26]. We did not observe any evidence for coding region hypermethylation in *rdr2* mutants. Instead, there was a clear reduction in non-CG methylation 5′ of the *ROS1* TSS in *rdr2* plants, as typically occurs when RdDM activity is lost (Fig. 3, S3 Fig.). Thus the TE at the 5′ end of *ROS1* is the most likely candidate as the site of RdDM activity that promotes *ROS1* expression, despite the fact that methylated TEs 5′ of genes are typically associated with transcriptional repression [18,20,21]. Combined with the distinct behavior of *ROS1* alleles inherited from *met1* or *rdr2* parents (Fig. 2), we propose that methylation by the RdDM pathway acts to promote *ROS1* expression via a different mechanism than does the MET1 pathway.

### Methylation 5′ of *ROS1* is sufficient to restore *ROS1* expression in *rdr2* mutants

Although our results suggest that *ROS1* expression is inversely correlated with DNA methylation at the *ROS1* locus, reduced expression of *ROS1* in RdDM mutants could be direct or indirect, for example due to altered expression of a *ROS1* regulator. To distinguish between these possibilities we sought to restore DNA methylation at the *ROS1* 5′ region in an *rdr2* mutant background and then observe the effect on *ROS1* expression. We attempted to bypass the inability of the *rdr2* mutant to make a dsRNA that initiates RdDM by expressing an inverted repeat transgene under the control of the constitutive 35S promoter in *rdr2* mutants (Fig. 4A). Transcription of inverted repeat transgenes creates a double-stranded hairpin RNA that can be processed into small RNAs that direct DNA methylation [31]. We used inverted repeats corresponding precisely to the 228 bp *ROS1* 5′ region that is methylated in wild type. We screened *rdr2* T_1_ plants for methylation of the *ROS1* 5′ region and focused on six lines for in-depth analysis of DNA methylation by bisulfite sequencing (Fig. 4B). Methylation in each of the six lines was restored to varying degrees, constituting an epiallelic series, with some lines exhibiting a methylation profile strikingly similar to wild type. Methylation occurred in all sequence contexts, indicative of RNA-directed DNA methylation. *ROS1* expression was examined in leaves of each independent line by RT-qPCR (Fig. 4C). Remarkably, *ROS1* expression was restored in *rdr2* mutants when methylation of the 5′ region was restored. In a line with limited restoration of DNA methylation (line #19), *ROS1* expression increased only marginally in *rdr2* mutants (Fig. 4C). These data demonstrate that methylation of the 5′ sequence is sufficient to promote *ROS1* expression, and eliminate the possibility that decreased expression of *ROS1* in *rdr2* mutants is caused by an indirect mechanism.

**Fig. 4:**
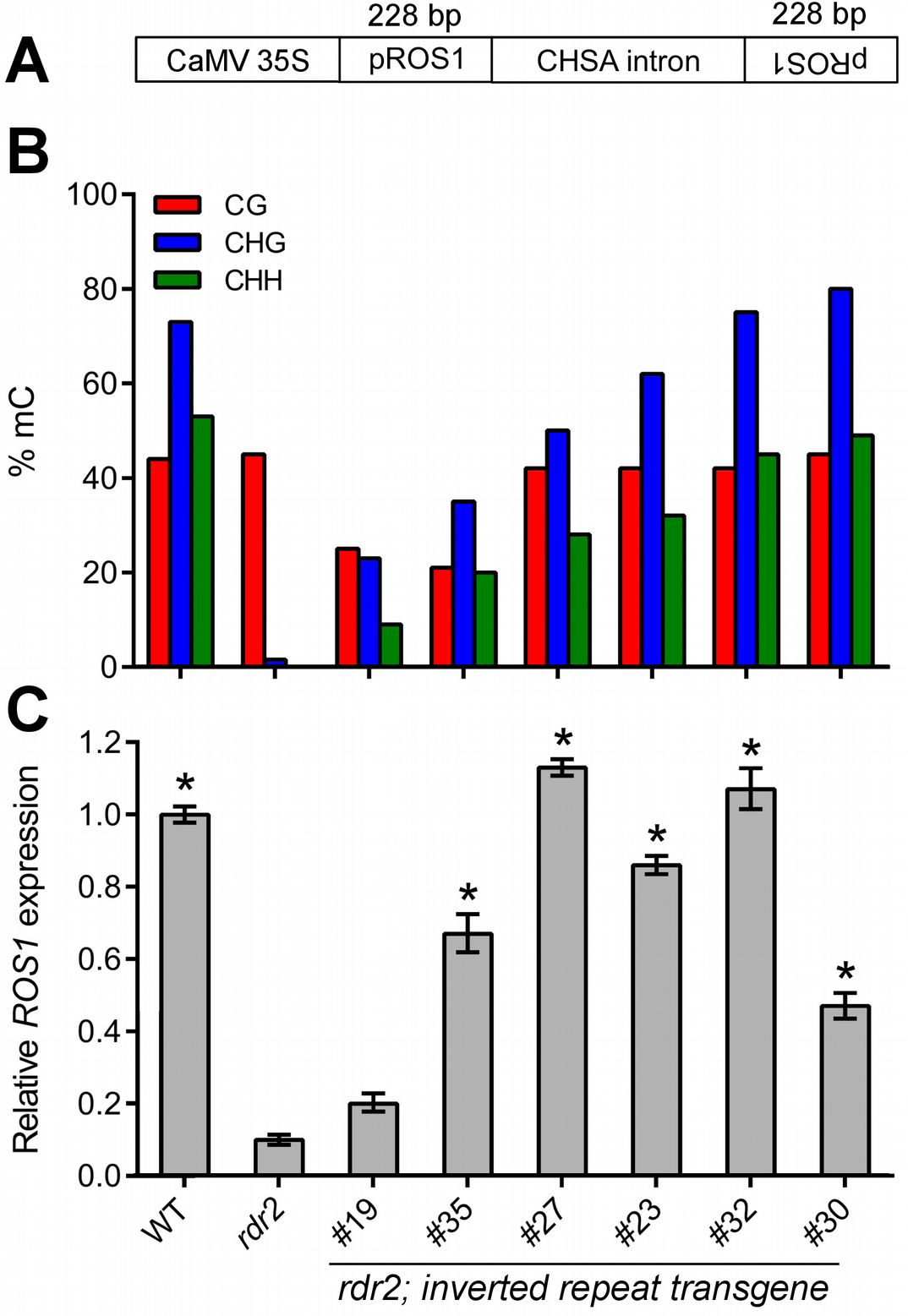
Restoration of methylation 5′ of *ROS1* is sufficient to restore *ROS1* expression in *rdr2* mutants. **(A)** The inverted repeat construct used to express double stranded RNA matching *ROS1* in *rdr2* mutants. CaMV 35S: Cauliflower mosaic virus 35S promoter; CHSA: Chalcone synthase. **(B)** Methylation from -595 to -534 bp before the translational start site of *ROS1* in WT (Col-0), *rdr2* mutants and six independent transgenic lines expressing the inverted repeat transgene in an *rdr2* background. **(C)** Quantification of *ROS1* expression in WT, *rdr2* and each of the transgenic lines. *p = <0.001 compared to *rdr2,* two-tailed t-test.

### ROS1 directly opposes RdDM to repress *ROS1* expression

We noticed that cytosines in the CG context 5′ of *ROS1* were intermediately methylated at around 50%, in wild-type tissues with independent variance at every CG position in the sequence (Fig. 3, S3 Fig.). CG methylation is faithfully copied by MET1 during DNA replication, and so average methylation at symmetric CG sites is usually close to 0 or 100% [20]. The observed intermediate level of CG methylation and the low frequency bisulfite clones fully methylated in the CG context (S3 Fig.) suggested that DNA methylation and DNA demethylation might both be active at the 5′ sequence. We hypothesized that ROS1 might oppose RdDM to remove methylation at its own promoter. We performed bisulfite sequencing of the 5′ methylated region in two missense mutants of *ROS1, ros1-2* [32] and *ros1-7,* an allele encoding a protein with an E956K substitution in the ROS1 DNA glycosylase domain [33]. Symmetric CG methylation increased to nearly 100% in both mutants compared to their wild-type siblings, along with increases in non-CG methylation (Fig. 5A), indicating that ROS1 actively removes methylation from this region in wild-type plants. At other loci, removal of 5′ methylation by ROS1 increases transcription [21]. We examined the effect of DNA demethylation by ROS1 on *ROS1* transcription by performing RT-qPCR on *ros1-2* and *ros1-7* mutants. Because these are missense mutations, nonsense-mediated decay should not be a complicating factor in measuring transcript abundance. *ROS1* transcripts were 2 to 4-fold more abundant in *ros1* mutants (Fig. 5B). This suggests that active demethylation by ROS1 represses transcription of *ROS1,* counteracting the function of the RdDM pathway, which promotes *ROS1* expression. Thus ROS1 regulates the expression of its own gene, forming a negative feedback loop for demethylation activity.

**Fig. 5:**
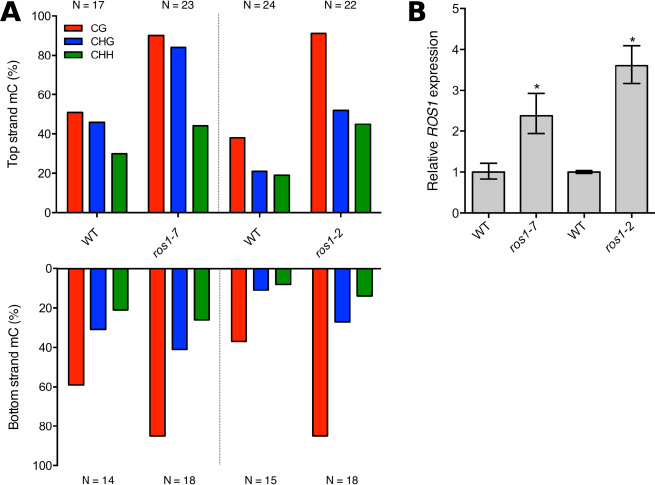
ROS1 demethylates the *ROS1* promoter and reduces expression. **(A)** Methylation of the 5′ region of *ROS1* (region 1 in Fig. 2) was measured in two *ros1* missense mutants by bisulfite PCR. WT strains were segregating Col-0 siblings for *ros1-7* and were C24 for *rosl-2*. N represents the number of independent clones sequenced for each BS-PCR product. **(B)** Relative *ROS1* transcript abundance was measured in seedlings of both mutants using RT-qPCR. Data represented as mean, error bars represent standard deviation. *p = <0.005, two-tailed t-test.

### Methylation-sensitive regulation of *ROS1* is conserved in other species

To determine if regulation of *ROS1* by methylation might be adaptive, we assessed whether methylation-sensitive expression of *ROS1* is conserved in other species. *Arabidopsis lyrata,* which diverged approximately 10 million years ago, has two highly conserved paralogs of *ROS1* in tandem in the genome, which we termed *AlROS1a* and *AlROS1b* (Fig. 6A). We performed a Bayesian reconstruction of the phylogeny of *ROS1* homologs within all sequenced Brassicales (Fig. 6B) and found that the duplication giving rise to two *ROS1* paralogs in *A. lyrata* occurred prior to the divergence of *A. lyrata* from *A. thaliana. AtROS1* belongs to the same clade as *AlROS1a,* and no true homologs to *AlROS1b* exist in *A. thaliana* (Fig. 6B). The homolog to *AlROS1b* was likely lost in the lineage that gave rise to *A. thaliana. AlROS1a* and *AlROS1b* share a high degree of sequence similarity in their coding region, but no significant similarity in their upstream sequences. Only *AlROS1a* has an upstream region conserved with *AtROS1* (Fig. 6A), including the presence of the same 5′ TE. The 5′ sequences are 78% identical over the first 1.4 kb.

**Fig. 6:**
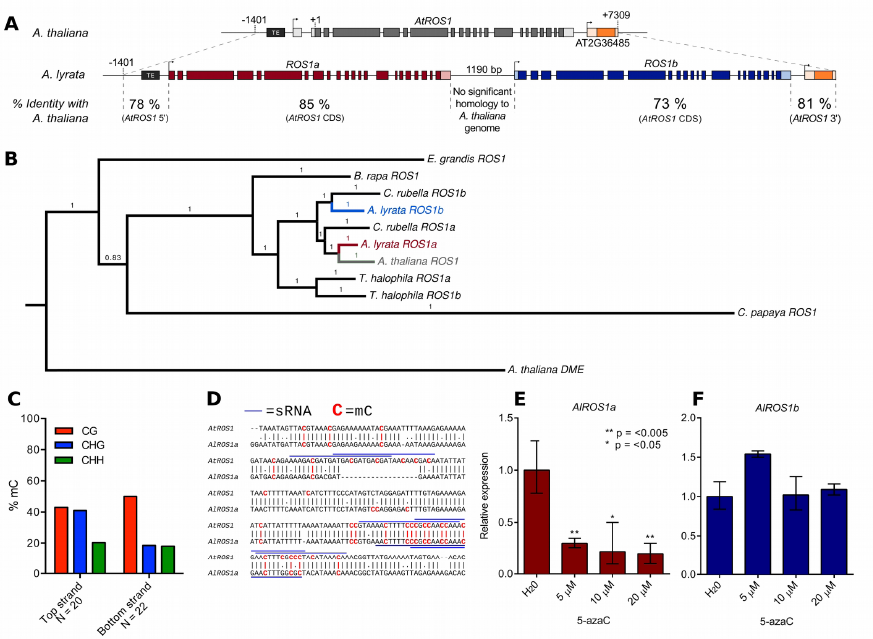
Methylation-sensitive expression of *ROS1* is evolutionarily conserved. **(A)** *ROS1* is locally duplicated in the closely related species *Arabidopsis lyrata*. Percentage DNA sequence identity between each paralog and *A. thaliana ROS1* is shown for upstream, coding and downstream regions. **(B)** Bayesian reconstruction of the phylogeny of *ROS1* homologs within Brassicales shows *AIROSIa* and *AIROSIb* diverged prior to the divergence of *A. thaliana* and *A. lyrata*. Support values are Bayesian posterior probabilities. (C) Methylation of the sequence 5′ of *AIROSIa* was examined by BS-PCR. N represents the number of independent clones sequenced for each BS-PCR product. **(D)** Annotation of methylated cytosines in the *AtROSI* and *AIROSIa* 5′ region. Blue lines annotate sequences matching sRNAs from their respective species. **(E)** Relative *AIROSIa* transcripts are significantly decreased when plants are grown on 5-azaC. **(F)** Relative *AIROSIb* transcripts levels are unaffected by growth on 5-azaC. Data represented as mean, error bars represent standard deviation. *p = <0.05, **p = <0.005, two-tailed t-test.

We performed bisulfite sequencing of the *AlROS1a* 5′ region (Fig. 6C) and of the exonic sequences in *AlROS1a* and *AlROS1b* that match exons 15-18 in *ROS1* (S4 Fig.). Although CG methylation was present in the 3′ exonic region of both paralogs, non-CG methylation was absent (S4 Fig.). Additionally, unlike *A. thaliana,* we did not find any small RNAs matching these exons for either *AlROS1a* or *AlROS1b* in a small RNA dataset from *A. lyrata* flowers [34]. By contrast, the methylation profile 5′ of *AlROS1a* was remarkably similar to the methylation profile 5′ of *AtROS1* (Fig. 6C-D and Fig. 3). Like *AtROS1*, CG methylation in the 5′ region was at an intermediate level between 40-50%, suggesting that RdDM and active DNA demethylation might also simultaneously target *AlROS1a*. Pair-wise alignment of the sequences from each species showed that the methylation is present at conserved cytosines (Fig. 6D). Furthermore, we identified small RNAs from *A. lyrata* datasets [34] that are almost identical to the small RNAs associated with methylation at the 5′ end of *AtROS1* (Fig. 6D). The methylated sequence matching these RNAs is directly adjacent to a homolog of the TE upstream of *AtROS1* (Fig. 6A). These data indicate that RdDM targeting to a region 5′ of the *ROS1* TSS is evolutionarily conserved.

To test whether the expression of either *AlROS1* paralog was responsive to methylation alterations, *A. lyrata* seedlings were grown on varying concentrations of 5-azaC. *AlROS1a* but not *AlROS1b* transcripts were significantly decreased in seedlings grown on 5-azaC (Fig. 6E-F). Thus expression of the true *A. lyrata* homolog of *ROS1, AlROS1a,* which has similar 5′ methylation and conserved small RNAs, is methylation-responsive. Together, these data suggest that the regulation of *ROS1* by RdDM and DNA demethylation at 5′ sequences is conserved between *A. thaliana* and *A. lyrata*. Interestingly, in transcriptome datasets from shoot apical meristems or immature ears of *Z. mays mop1* mutants *(Mop1* is a *RDR2* homologue), expression of two DNA glycosylase genes with high homology to *AtROS1 (DNG101* and *DNG103)* is reduced 2 to 3.3-fold in comparison to wild type [35,36]. Reduced expression of ROS1 homologs has also been observed in the transcriptome of *Z. mays* RNA polymerase 4 mutants [37]. We confirmed that DNA glycosylase expression is reduced in *mop1* leaves by RT-qpCR (S5 Fig.). Both *DNG101* and *DNG103* have methylated TEs in their 5′ region in all sequence contexts, suggesting that both loci could be direct targets of RdDM [38]. These data further suggest that regulation of DNA glycosylases by RdDM might be a general feature of angiosperms, and thus likely adaptive.

## DISCUSSION

Our data demonstrate that *ROS1* functions as a self-regulating epigenetic rheostat. RdDM and DNA demethylation activities converge at the *ROS1* locus, but each activity has the opposite outcome on *ROS1* transcript abundance compared to typical targets of these processes (Fig. 7). By establishing DNA methylation at the *ROS1* locus in *rdr2* mutants (Fig. 4), we have conclusively shown that methylation of the 5′ sequence is sufficient to restore *ROS1* expression. Thus reduced expression of *ROS1* in *rdr2* mutants is not caused by an indirect mechanism, such as decreased expression of another gene required for *ROS1* expression or increased expression of a negative regulator of *ROS1*. While the precise mechanism by which the activity of *ROS1* is repressive and RdDM is activating remains unknown, we speculate that a protein that either negatively or positively regulates *ROS1* may exhibit differential DNA binding based on methylation of the underlying *ROS1* 5′ DNA sequence. Alternatively, it is possible that rather than DNA methylation itself, the act of RdDM could play a regulatory role. For example, occupancy, DNA melting or elongation by the RNA polymerases IV or V could be required for positive regulatory factors to access *ROS1*. Interestingly, RdDM in an intron of the *MADS3* gene in Petunia has also been shown to be associated with transcriptional upregulation [39]. In this instance, it is likely that methylation of a short cis-element is necessary to confer increased expression.

**Fig. 7:**
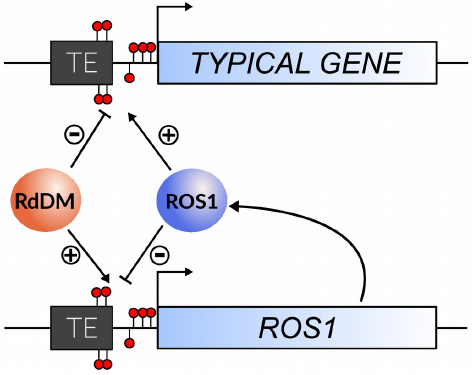
The regulation of *ROS1* by methylation acts as an epigenetic rheostat. A schematic summarizing the roles of RdDM, and ROS1 at the *ROS1* locus and other genomic targets. Demethylation by ROS1 counter-acts RdDM to reduce *ROS1* expression, in opposition to its function at typical targets. This ensures a balance of methylation and demethylation activities is maintained in the genome.

The reversal of methylation outcomes at *ROS1* permits the genome to maintain gene expression (promoted by DNA demethylation) and genome defense (TE silencing by RdDM) in homeostatic balance. For example, if the genome were under stress from high TE transcriptional activity or invasion, RdDM activity might increase. Under these conditions, RdDM activity would promote the expression of *ROS1,* so that the activity of demethylation at target genes would be maintained in equilibrium with the activity of RdDM in the genome (Fig. 7). Conversely, if RdDM were less active, *ROS1* expression would be reduced to prevent hypomethylation as a consequence of demethylation activity. The result is that the expression of *ROS1* is always maintained in balance by its autoregulation (Fig. 7), which may help underpin the regulation of epigenetic homeostasis within plants and explain why spontaneous changes to methylation are generally very rare [40,41]. This homeostatic balance may be dynamically modified in certain conditions, such as fungal or bacterial infections, where active demethylation of defense genes is important for resistance [2–4]. The rheostat may also be important during normal development. In pollen, *DRM2* is expressed at a low level in microspores and sperm [8]. In agreement with our findings from *drm2* and other RdDM mutants, *ROS1* transcripts have not been detected in sperm [42]. Published methylation profiling data from pollen show that the *ROS1* 5′ region is hypomethylated in sperm cells, but not in the vegetative nucleus (S6 Fig.), where both *DRM2* and *ROS1* are expressed [42,43]. In the future, it will be important to determine whether disrupting the rheostat has consequences for the proper establishment of methylation patterns in sperm or in developing progeny after fertilization.

In addition to the methylation-responsive regulation of *ROS1* that we have described here, it is possible that additional mechanisms underlie epigenetic homeostasis in *Arabidopsis*. Targeting of RdDM, H3K9 methylation and H3K27 methylation are redirected throughout the genome in *met1* mutants [20,24,44], and it has been hypothesized that this redirection may be the result of compensatory mechanisms necessary to maintain some level of integrity in gene expression [24]. Other mechanisms may also therefore regulate repressive histone modifications in balance with gene expression. The role of methylation in promoting *IBM1* expression [26] may be one such example. Although our experiments have focused on Arabidopsis species, the concept of epigenetic homeostasis might also apply more broadly to other DNA methylation systems, including those in mammals. It is known that the cancer epigenome exhibits global DNA hypomethylation and local hypermethylation [45], which is broadly similar to the methylation phenotype of a *met1* mutant. Interestingly, expression of the three *TET* enzymes, which are responsible for initiating DNA demethylation in mammals by oxidizing 5-methylcytosine, is reduced in multiple cancers [46].

We conclude that the *ROS1* locus serves as a rheostat for methylation levels. We propose that the *ROS1* epigenetic rheostat evolved to counter-balance positive feedback between DNA methylation and RdDM activity [13,14] to prevent ectopic gain of DNA methylation. The conservation of methylation-sensitive *ROS1* expression among divergent angiosperms suggests that this regulation is adaptive and could underpin how plants balance a number extremely effective, potent, and self-reinforcing silencing mechanisms while maintaining gene transcription.

## MATERIALS and METHODS

### Plant growth conditions

*Arabidopsis thaliana, Arabidopsis lyrata,* and *Zea mays* plants were grown in a greenhouse with 16-hour days at 21°C. For experiments performed on whole seedlings, plants were grown on 0.5 x MS medium with 5% agar. For treatment with 5-azaC, *A. thaliana or A. lyrata* seedlings were grown on filter paper moistened with water or 5, 10, 15 or 20 μM 5-aza-2-deoxycytidine. Fresh water or 5-azaC was added daily. The accession number for all mutant plants used in this study are in the Supplemental Materials and Methods.

### Quantitative RT-PCR

Total RNA was extracted using a Plant RNA extraction Kit (Qiagen). RNA was extracted from whole 7-day old *Arabidopsis thaliana* seedlings for all experiments, except for experiments using 5-azaC, in which case 5-day old *A. thaliana* or *A. lyrata* seedlings were used, or experiments with transgenic lines expressing a *ROS1* inverted repeat construct, in which case rosette leaves from 21-day old plants were used. RNA was extracted from the tip of the third true leaf of maize plants. For each genotype, 3 biological replicates of at least 5 pooled individual seedlings *(Arabidopsis)* or individual plants (maize) were collected. Genomic DNA was removed using amplification-grade DNAseI (Invitrogen). cDNA was synthesized from 500 ng RNA using Superscript II reverse transcriptase (Invitrogen) according to manufacturers′ instructions, selecting for polyadenylated transcripts using an oligo-dT primer. For each cDNA synthesis reaction, a control was performed without addition of reverse transcriptase to test the efficacy of the DNAse treatment. Quantitative RT-PCR (RT-qPCR) was performed using Fast Sybr-Green mix (Applied Biosystems) according to manufacturers′ instructions. All reactions were performed using a StepOne Plus Real-Time PCR system (Applied Biosystems). Primers were designed to have matching melting temperatures between 60-65°C and to produce amplicons between 80-160 bp in length. All primers were used in a final concentration of 400 nM. The efficiency of all primer pairs was verified using a standard curve dilution of template cDNA prior to their use in quantification of transcripts. Melt curves were analyzed to verify the presence of one amplicon in each reaction, and representative products were also verified by agarose gel electrophoresis. Relative expression was calculated using the ddCt method as described [47]. For *Arabidopsis,* the reference transcript used for all reactions was AT1G58050, experimentally verified to be one of the most consistently abundant transcripts in *A. thaliana* [48]. For maize, the reference transcript was *ZmEF1α*, defined to be the most consistent reference transcript over the majority of experimental conditions [49]. Primer sequences are available in the Supplemental Materials and Methods.

### Rapid Amplification of cDNA ends (RACE)

5′ RACE of *ROS1* was performed using 10 μg Col-0 RNA extracted from 10-day old seedlings. The 5′ RACE cDNA library was synthesized using a FIRST-CHOICE RLM-RACE Kit (Ambion) according to manufacturers instructions, with the exception that a ROS1-specific oligonucleotide was used to prime cDNA synthesis. RACE products were amplified using a nested PCR strategy, purified using a Qiaquick gel extraction kit (Qiagen) and cloned for sequencing using a TOPO-Blunt PCR cloning kit (Invitrogen).

### Bisulfite sequencing

Genomic DNA was isolated from 7-day old seedlings or 21-day old rosette leaves using a CTAB procedure. 2 μg DNA were sheared by sonication and used for bisulfite treatment, which was performed as described [50]. 2 μl bisulfite treated DNA was used in PCR reactions with 2.5 U ExTaq DNA polymerase (Takara) and 0.4 μM primers using the following cycling conditions (95°C 3 minutes, 40 cycles of [95 °C for 15 seconds, 52 °C for 60 seconds, 72 °C for 60 seconds], 72 °C for 10 minutes). PCR products were cloned using TOPO-TA PCR cloning kit (Invitrogen) and individual colonies were sequenced. Sequenced products were aligned using MUSCLE [51], and methylation of each cytosine residue was calculated using CyMate [52].

### Inverted repeat transgene

The 228 bp sequence 5′ of *ROS1* that is targeted by RdDM was amplified and cloned into the directional entry vector pENTR-TOPO-D (Invitrogen). The sequence was then inserted twice in an inverted repeat conformation into the vector pANDA-35HK using a single LR clonase reaction as described by [53]. *rdr2* mutant plants were transformed with the inverted repeat transgene by floral dipping [54], and T_1_ lines were screened for DNA methylation 5′ of *ROS1* using a restriction enzyme assay on bisulfite treated DNA. 90% of lines screened exhibited higher DNA methylation than *rdr2* and *nrpe1* mutants. Six lines covering a range of methylated epigenotypes were selected for bisulfite sequencing and expression analysis.

### Bayesian reconstruction of phylogeny

Coding sequences for *ROS1* homologs from all fully sequenced genomes within Brassicales were downloaded from Phytozome 9.1. In addition, the coding sequence for *Arabidopsis thaliana DME,* which belongs to a distinct clade of DNA glycosylases [55], was included as an out-group. Sequences were aligned using Muscle 3.8 [51], and manually verified using Aliview [56]. The phylogeny was reconstructed from the aligned matrix using MrBayes 3.1.2 [57]. jModelTest [58] was used to determine the ideal model for analysis, and the general time reversible model with gamma distributed rate variation was chosen. Gaps were treated as missing data. The analysis was run for 2,500,000 generations sampling every 100 trees. By this time the average standard deviation of split frequencies had reached <0.01 and the potential scale reduction factor (PRSF) was <1.005 for all parameters. The first 25% of trees were discarded as burn-in and the output phylogeny was further analyzed and annotated using FigTree 1.4.

## Acknowledgements

We thank Jennifer Cooper and Christine Codomo for isolating the *ros1-7* allele from the Arabidopsis TILLING population, Nathan Springer for *mop1* maize seed, Jian-Kang Zhu for *ros1-2* seed, and Eric Richards for *vim1; vim2; vim3* seed. MG is a Pew Scholar in the Biomedical Sciences and an Alexander and Margaret Stewart Trust Scholar.

## Supplementary Material

**S1 Figure:**
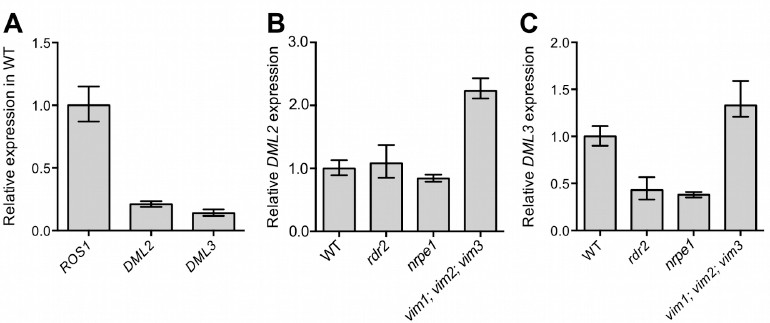
*DML2* and *DML3* expression in methylation mutants. **(A)** The quantification of *DML2* and *DML3* transcripts in WT (Col-0) by RT-qPCR, normalized to the abundance of *ROS1* transcripts. **(B)** *DML2* expression and **(C)** *DML3* expression in RdDM and maintenance methylation mutants. Data represented as mean, error bars represent standard deviation.

**S2 Figure:**
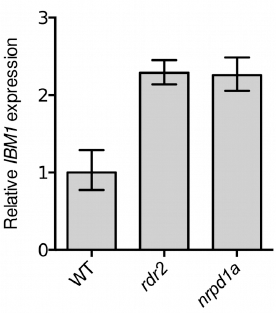
*IBM1* is not silenced in RdDM mutants. The level of expression of the long transcript isoform of *IBM1* was measured using seedlings of two RdDM mutants using RT-qPCR. In both *rdr2* and *nrpd1a* mutants, IBM1 expression was higher than in WT (Col-0) at p = <0.05. Data represented as mean, error bars represent standard deviation.

**S3 Figure:**
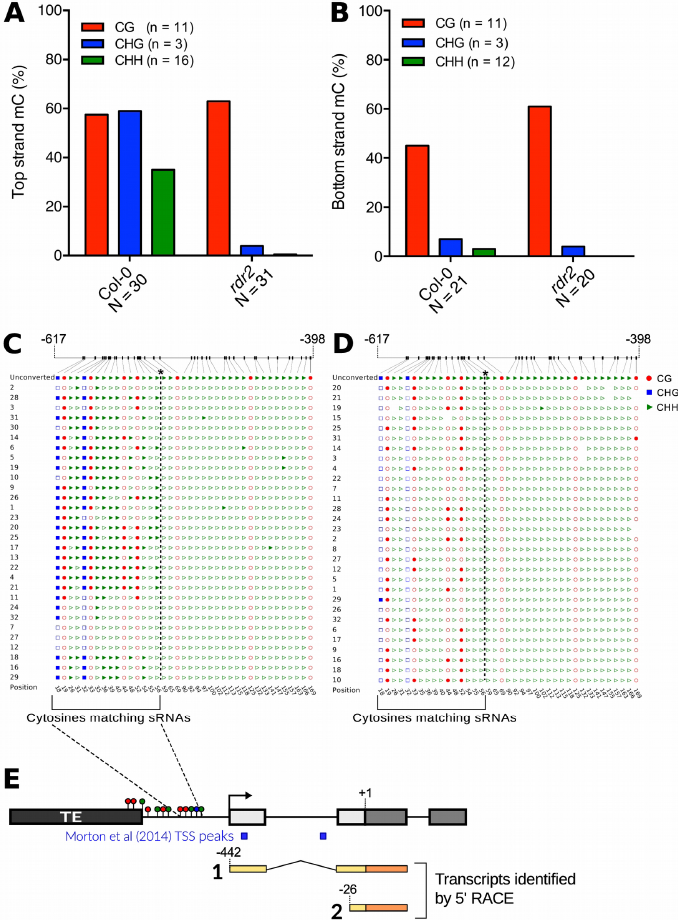
The *ROS1* promoter is heavily methylated by RdDM. **(A-B)** Methylation levels of the top and bottom strand of a 200 bp region upstream of the *ROS1* transcription start site were quantified using BS-PCR on DNA from wild-type Col-0 and *rdr2* mutants. Percentage methylation was calculated from the first to last methylated cytosine. N represents the number of independent clones sequenced; n represents the number of cytosines counted in the region. **(C)** and **(D)** show a portion of the cytosines analyzed from the top strand of Col-0 and *rdr2,* respectively. Filled/unfilled shapes represent methylated/unmethylated cytosines respectively. *The dotted line indicates the 3′-most position from which methylation was quantified. The quantified region continues beyond what is shown at the 5′ end **(E)** The transcription start sites of *ROS1* in WT were determined using 5′ RACE. The position of start sites identified by RACE and in a genome-wide profiling study [63] are shown relative to the region of DNA methylation.

**S4 Figure:**
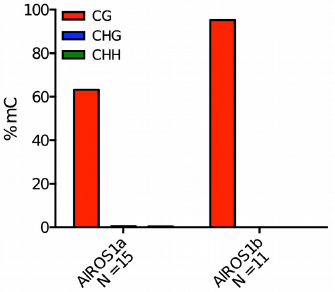
*AIROS1a* and *AIROS1b* share similar coding region methylation. DNA methylation in exons 19-20 of *AIROSIa* and exons 18-19 *AIROSIb* (which share homology with *AtROS1* exons 17-18) was measured using BS-PCR from leaf DNA. Percentage methylation in each sequence context was calculated from the first to the last methylated cytosine. 6 GC, 2CHG and 32 CHH sites were counted for *AIROSIa*. 7 CG, 11 CHG and 31 CHH sites were counted for *AIROSIb.*

**S5 Figure:**
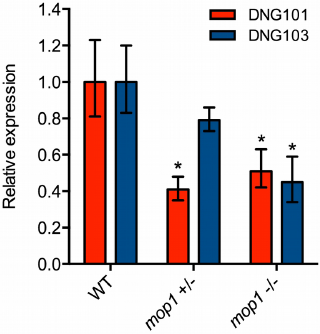
Two *ROS1* homologs are down-regulated in a maize RdDM mutant. Transcript abundance of the 5-methylcytosine DNA glycosylases *DNG101* and *DNG103* was measured in maize plants segregating for a mutation in *Mop1,* a homolog of RdDM, using RT-qPCR. WT is the inbred strain B73. Data are represented as mean, error bars represent standard deviation. *p = <0.01, two-tailed t-test.

**S6 Figure:**
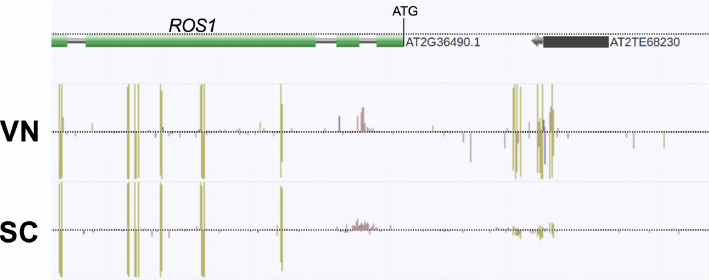
*ROS1* is hypomethylated in sperm cells. A snapshot of genome-wide methylation data from Arabidopsis pollen [42], showing the 5′ region of *ROS1*. CG, CHG and CHH methylation are denoted by green, blue and red lines, respectively. VN = vegetative nucleus. SC = sperm cells.

## Supplementary Materials & Methods

### Arabidopsis mutants

The mutants used for this study were obtained from the Arabidopsis Biological Resource Center (ABRC) [59], Ohio State University, unless otherwise stated.

*ago4:* accession CS9927

*ago4; ago6:* accession CS66095

*cmt2:* CS849188

*cmt3:* CS3665

*dcl3:* CS16390

*ddm1: ddm1-5* [60]

*dms3:* SALK_068723C

*drm1; drm2:* CS6366

*hda6:* CS66153 *(axe1-5)*

*ibm1:* SALK_035608C

*kyp:* SALK_044606C

*met1: met1-6* [50]

*nrpd1a:* SALK_128428

*nrpe1:* SALK_029919C

*rdr2-1:* CS66076

*rdr6:* CS24285

*ros1-2:* Obtained from Jian-Kang Zhu [32]

*ros1-7:* Obtained from the TILLING project [33]

*shh1:* SALK_074540C

*spt5l:* SALK_001254C

*vim1; vim2; vim3:* Obtained from Eric Richards [61]

*mop1-1 (maize):* Obtained from Nathan Springer [62]

### Primers used in this study

#### RT-qPCR

ROS1 F: CAGGCTTGCTTTTGGAAAGGGTACG

ROS1 R: GTGCTCTCTCACTCTTAACCATAAGCT

DML2 F: CGGGAAGAGGAATCACAGACT

DML2 R: GGACGTCGATAGGGTTTATGCT

DML3 F: CGTAGGGAGTTGTGTAAGGGA

DML3 R: GCAAAGTTCAATCCGTCTTGTGT

IBM1 F: TGCTGTCCTGTGTCTCAGGTTG

IBM1 R: ACCGCGTCAGATAGAAGTTCTGG

AT1G58050 F: CCATTCTACTTTTTGGCGGCT

AT1G58050 R: TCAATGGTAACTGATCCACTCTGATG

AlROS1a/b F: TTGCTATTTGGACGCCAGGTGAG

AlROS1a R: ATACACTTACTTACAGCCGGTTG

AlROS1b R: ATACACTTGCTCACAGTTGGTTG

Al315392* F: CCATTCTACTTTTTGGCGGCT

Al315392* R: TCAATGGTTACTGATCCACTCTGATG

*\*A. lyrata* homolog to AT1G58050

*DNG101* F: CCAGATGATCCCTGTCCATATCTTC

*DNG101* R: GGCATCGATCGATTGTGCAGTTTC

*DNG103* F: CCATGCTGTGACCCTCAAATG

*DNG103* R: CTCTGCAGTACAATTGTGGCAC

*ZmEF1α* F: TGGGCCTACTGGTCTTACTACTGA

*ZmEF1α* R: ACATACCCACGCTTCAGATCCT

#### Bisulfite Sequencing

*AtROS1* 5′ end top strand 1 F: GAYTAAAAYATTTGGAATGATYAAAAAYGAAAG

*AtROS1* 5′ end top strand 1 R: TTGTTTTCTACAAAATCTCCTARACTAT

*AtROS1* 5′ end bottom strand 1 F: CAACTARCCTAATAATCACTCTACTACACT

*AtROS1* 5′ end bottom strand 1 R: TAGAYTATGGGAAAGATGATTTAAAAAG

*AtROS1* 5′ end top strand 2 F: GGAGATTTTGTAGAAAAGAATYATT

*AtROS1* 5′ end top strand 2 R: TCACTRATRCTTCRTTTCTTCTCTT

*AtROS1* 5′ end bottom strand 2 F: CTTTTTAAATCATCTTTCCCATARTCTA

*AtROS1* 5′ end bottom strand 2 R: GTAGAATYAATGGTTATGGTGGTG

*AtROS1* coding region fragment 2a F: CACAAACCTTTCCTCCAATTRACTRCTAT

*AtROS1* coding region fragment 2a R: GATTYYAAGAGAAGAAAYGAAGYATYAGTG

*AtROS1* coding region fragment 2b F: CACAAACCTTTCCTCCAATTRACTRCTAT

*AtROS1* coding region fragment 2b R: GATYTGYYYATAYAYGGTGGAGGAT

*AtROS1* coding region fragment 3 F: CATCCCRCRACTCTTRATTRTTTCARCAAC

*AtROS1* coding region fragment 3 R: GAGYAGGATYAAGYTYAGAGATYGAYTTAG

*AtROS1* coding region fragment 4 F: CRCCTCCTCTATRTCARCTATTRATACTTC

*AtROS1* coding region fragment 4 R: GYYAGTGYGTTTGYAAGGTGYTAYAAYATG

*AtROS1* coding region fragment 5a F: TTGTGGYAGTTGGAAAAGAGAGAAYYTG

*AtROS1* coding region fragment 5a R: CTTACCTCATTTACTTRAAARTACRTTCC

*AtROS1* coding region fragment 5b F: GGAAYGTAYTTTYAAGTAAATGAGGTAAG

*AtROS1* coding region fragment 5b R: CCACRTACACATACRTACCCTACATA

*AlROS1a* 5′ end top strand F: TGTTAAAGAAAAGGATAGAAYATGTGTG

*AlROS1a* 5′ end top strand R: CCTAATAATCACCCTATAACTTCCT

*AlROS1a* 5′ end bottom strand F: TCTTTCTCTAACTTTCATARCCRTTT

*AlROS1a* 5′ end bottom strand R: AAGAATTGTAAGGGGAYTAGYYTAAT

*AlROS1a* exons 19-20 F: TTGYTAGTGYGTTTGYAAGGTGYTA

*AlROS1a* exons 19-20 R: CATCCTCTATRTCARCTATTRATACTT

*AlROS1b* exons 18-19 F: GTATGTGAAGTAAYTGGATAAGGAYAYATG

*AlROS1b* exons 18-19 R: CTTACCTCATTTACTTRAAARTACRTTCC

### 5′ RACE

*ROS1* cDNA synthesis primer: CTCACAGTCACCCGCGTATCA

*ROS1* RACE outer nested PCR: GACTGCTATGATATTGATCCTCCAC

*ROS1* RACE inner nested PCR: GCTTTCTTCTCTCCTCTGTTTCTCCAT

### Transgene construction

Inverted repeat F: CACCGTTAGTTCATATAATTTTAAATAGTTACGT

Inverted repeat R: AGGGCGAAAGTTCGTTTGGTTG

